# Microglia from offspring of dams with allergic asthma exhibit epigenomic alterations in genes dysregulated in autism

**DOI:** 10.1101/192997

**Authors:** Annie Vogel Ciernia, Milo Careaga, Janine LaSalle, Paul Ashwood

## Abstract

Dysregulation in immune responses during pregnancy increase the risk of a having a child with an autism spectrum disorder (ASD). Asthma is one of the most common chronic diseases among pregnant women, and symptoms often worsen during pregnancy. We recently developed a mouse model of maternal allergic asthma (MAA) that induces changes in sociability, repetitive and perseverative behaviors in the offspring. Since epigenetic changes help a static genome adapt to the maternal environment, activation of the immune system may epigenetically alter fetal microglia, the brain’s resident immune cells. We therefore tested the hypothesis that epigenomic alterations to microglia may be involved in behavioral abnormalities observed in MAA offspring. We used the genome-wide approaches of whole genome bisulfite sequencing to examine DNA methylation and RNA sequencing to examine gene expression in microglia from juvenile MAA offspring. Differentially methylated regions (DMRs) were enriched for immune signaling pathways and important microglial developmental transcription factor binding motifs. Differential expression analysis identified genes involved in controlling microglial sensitivity to the environment and shaping neuronal connections in the developing brain. Differentially expressed associated genes significantly overlapped genes with altered expression in human ASD cortex, supporting a role for microglia in the pathogenesis of ASD.

**Main Points:** Maternal allergic asthma induces changes in DNA methylation and transcription in juvenile offspring microglia

Differentially methylated regions are enriched for functions and transcription factor binding motifs involved in inflammation and microglial development

Differentially expressed genes and differentially methylated regions are enriched for genes dysregulated in Autism Spectrum Disorders

## Introduction

Epigenetic mechanisms fine tune gene expression during the development and maturation of the nervous and immune systems (Vogel Ciernia and LaSalle, 2016) resulting in long-lived phenotypic alterations without changing DNA sequence. Epigenetic mechanisms such as DNA methylation can be altered by environmental changes during a particular critical window in early life (Dolinoy et al., 2007; Jirtle and Skinner, 2007). The developing mammalian brain is particularly sensitive to epigenetic alterations, as mutations in epigenetic effectors can result in human neurodevelopmental disorders (Lasalle, 2013; Vogel Ciernia and LaSalle, 2016). Epigenetic changes are thought to adapt a static genome to a dynamic environment, providing an important interface between genetic and environmental risk factors in the complex etiology of autism spectrum disorders (ASD).

Epidemiological reports suggest a strong association between periods of maternal immune activation (MIA) during pregnancy and an increased risk of having a child with ASD (Atladottir et al., 2009, 2010, Zerbo et al., 2013, 2015). Early case reports and small comparative studies focused on infections during pregnancy associated with ASD risk (Chess, 1977; Sweeten et al., 2004; Libbey et al., 2005). Large population based studies in Denmark showed that mothers with severe infections during pregnancy that required hospitalization were at increased risk of having a child with ASD (Atladottir et al., 2010); a finding also confirmed in a Californian cohort (Zerbo et al., 2013). Moreover, taking antipyretic medication to reduce fever during pregnancy eliminated this increased risk for ASD (Zerbo et al., 2013), suggesting that treating the MIA attenuates risk. In addition to infections, associations of risk for ASD with autoimmune and immunological conditions such allergy, asthma, celiac disease, type-1 diabetes, autoimmune thyroid disease, psoriasis, rheumatoid arthritis, and rheumatic fever have been reported (Croen et al., 2005; Mouridsen et al., 2007; Atladottir et al., 2009; Mostafa and Kitchener, 2009; Keil et al., 2010), providing additional evidence for maternal immune dysfunction and increased risk of ASD.

Asthma is one of the most common chronic diseases among pregnant women. During pregnancy the median percent of women hospitalized for acute exacerbations of asthma is 6% (Murphy et al., 2005, 2006, Ali and Ulrik, 2013a; b); however, in a study of 330 women with asthma, symptoms worsened in as many as 35% during pregnancy (Schatz et al., 1988). A large epidemiological study demonstrated that mothers with new onset allergies and asthma during the time of pregnancy were at increased risk for having a child with ASD (Croen et al., 2005). A separate study that looked at maternal allergies and asthma, irrespective of whether they were new onset or existing conditions, also found increased risk with certain allergies and ASD (Lyall et al., 2014). Recently we identified a profile of elevated serum cytokines midgestation in women who gave birth to a child later diagnosed with ASD (Goines et al., 2011; Jones et al., 2017). This cytokine profile of elevated interleukins IL-4 and IL-5 was consistent with an allergic asthma clinical phenotype (Magnan et al., 2000; Cho et al., 2005; Tamasi et al., 2005; Kumar et al., 2006). Furthermore, a different midgestation maternal cytokine profile of elevated IL-6 and IL-2 was associated with having a child with other developmental disorders, but not ASD (Goines et al., 2011). Amniotic fluid taken from children later diagnosed with ASD also had elevated levels of IL-4 whereas those with developmental delay had increased levels of IL-6 (Abdallah et al., 2012). These reports implicate asthma/allergy and associated cytokine responses as a potential contributing factor to the development of ASD.

Previous work in animal models examining the impact of gestational MIA on ASD-like behaviors of the offspring have focused on viral and bacterial responses, using polyinosinic-polycytidylic acid (polyI:C) or lipopolysaccharide (LPS) as experimental models (Whitehead et al., 2003; Nials and Uddin, 2008). However, allergies and asthma represent an alternative inflammatory response mediated by T-helper type 2 cells (T_H_2) that produce a different cytokine profile that may uniquely alter brain development relevant to ASD. We recently developed the first preclinical model of maternal allergic asthma (MAA) that produces the hallmark immunological responses in the MAA treated dams analogous to those observed in humans, including allergic airway inflammation and elevated levels of IL-4 and IL-5 (Schwartzer et al., 2015). MAA produced a phenotype in the offspring of impaired social approach and increased repetitive behaviors at both juvenile and adult ages (Schwartzer et al., 2015, 2017) that resemble behavioral features of ASD (Thomas et al., 2009). MAA challenge in gestation also resulted in altered brain neurochemistry in offspring similar to those observed in ASD (Schwartzer et al., 2015).

In humans, fetal exposure to maternal asthma led to epigenetic alterations in DNA methylation patterns in peripheral blood samples of the offspring (Gunawardhana et al., 2014). Notably, hypo-methylation in immune signaling gene *MAP8KIP3* in fetal blood correlated with blood eosinophils, nitric oxide levels, and total serum IgE in the mother (Gunawardhana et al., 2014), suggesting that human MAA responses may be related to additional epigenetic changes in the offspring. These findings are consistent with previous work which found histone acetylation (Tang et al., 2013) and DNA methylation (Basil et al., 2014; Richetto et al., 2016) changes in brain from juvenile and adult offspring in the polyI:C MIA mouse model. Together these results suggest that epigenetic changes within the immune cells of the brain may be responsible for the long-lived behavioral changes of offspring exposed to gestational MAA.

Within the brain microglia are brain-resident macrophages derived from erythro-myeloid precursors from the embryonic yolk sac that contribute to brain homeostasis and shaping synaptic and long-range neuronal connections (Tremblay et al., 2010; Paolicelli et al., 2011; Schafer et al., 2012; Hashimoto et al., 2013; Zhan et al., 2014). Microglia monitor and respond to their environment via specific cell surface receptors and secreted molecules, thereby acting as sentinel cells of tissue stress and injury (Murray and Wynn, 2011). The gestational timing of MAA may lead to alteration in microglial activity during key organizational phases of prenatal development producing long lasting changes in neural circuitry. Microglia arise from the yolk sac myeloid progenitors, migrate to the brain and colonize the neural folds during embryogenesis with minimal replacement from peripheral monocytes (Squarzoni, 2015), suggesting that any perturbations to microglia in early development may also persist through adulthood. To test the hypothesis that gestational MAA is associated with long-lived ASD-relevant epigenetic changes in microglia, we performed genome-wide analyses of the DNA methylome and transcriptome of microglia isolated from juvenile offspring. The results of this study provide genome-wide evidence that gestational MAA alters DNA methylation of key immune regulatory elements and transcription of genes dysregulated in human ASD brain, supporting the validity of the MAA mouse model, microglia as a relevant cell type, and maternal asthma as an ASD risk factor.

## Materials and Methods

### Animals

All experiments used C57Bl/6J mice from Jackson (Jax strain 000664). All experiments were conducted in accordance with the National Institutes of Health Guidelines for the Care and Use of Laboratory Animals. All procedures were approved by the Institutional Animal Care and Use Committee of the University of California, Davis.

### Maternal Allergic Asthma Paradigm

The MAA paradigm was conducted as previously described (Schwartzer et al., 2015, 2017). Briefly, on postnatal day (P) 42 and 49 sexually naive female C57Bl/6J mice were sensitized with a single intraperitoneal injection of 10 µg ovalbumin (OVA, Sigma, St. Louis, MO USA) in 1 mg (Al)OH_3_ (InvivoGen, San Diego, CA USA) dissolved in 200 µl phosphate buffered saline (PBS). One week following the second sensitization treatment, female mice were mated overnight and checked daily for the presence of a seminal plug, noted as gestational day 0.5 (G0.5). Pregnant mice were randomly assigned to either the allergic asthma or control group and exposed to either an aerosolized solution of 1% (wt/vl) OVA in PBS (MAA group) or vehicle control for three 45-minute induction on gestational days 9.5, 12.5, and 17.5, to correspond with early, middle, and late gestation. Following the final induction, mice were returned to their home cages, single housed, and left undisturbed until the birth of their litters. Pups remained with their mother until weaning on P21, at which time the offspring were group housed with same-sex littermates. For all experiments female offspring were chosen for analysis due to previously identified increased microglial reactivity at P30 in females compared to males (Schwarz et al., 2012). Female MAA offspring also show a similar behavioral phenotype to male MAA offspring with deficits in social interactions, increased marble burying and decreased grooming (Schwartzer et al., 2015, 2017).

### Microglial Isolation P35

P35 female mice were deeply anaesthetized with CO_2_ and then quickly perfused intracardially with ice-cold PBS. Whole brains were removed and stored on ice in HBSS without Ca^2+^/Mg^2+^ until processed. Each brain was gently homogenized to a single cell solution using a dounce homogenizer, then added to 1.8 ml of isotonic Percoll to form a 30% isotonic Percoll solution. Isotonic Percoll gradients were then constructed with layers of 70% with 1x phenol red, 37% and 30%. The resulting gradients were centrifuged for 30 min at 500xg at room temperature. The top layer of myelin and non-microglial cells were discarded and the microglia at the interface between the 37% and 70% layers were collected. The resulting microglia were then washed in HBSS and counted. The collected cells were immediately flash frozen for RNA/DNA extraction. Given the low yield from individual animals, an additional animal was run in parallel and harvested microglia were used to assess purity by flow cytometry for CD45.2 and CD11b. Gating was first performed on live single cells (96%). Any highly granular myelin contamination was removed (92% remaining). From that >90% was microglia (CD45.2 and CD11b positive). The remaining ≈6% (CD45.2 positive, but CD11b negative) represent a minimal contamination of other mononuclear cells (Figure S1).

### RNA sequencing library preparation and sequencing

RNA and DNA were isolated from the same isolated microglia samples using ZR-Duet RNA/DNA kit (Zymo, D7001) with on-column DNase digestion (Qiagen, 79254) for RNA extraction. RNA and DNA were quantified using Qubit High Sensitivity assays. RNA quality was assessed by Bioanalyzer and only samples with RNA Integrity Number (RIN) scores greater than 7 were used for library preparation. 60-100 ng of input RNA per sample were used for RiboGone rRNA removal (Clonetech, 634847). 2 ng of rRNA depleted RNA per sample was then used for library construction using the SMARTer Stranded RNA-Seq kit (Clontech, 634836) and 14 cycles of PCR amplification. Library quality was assessed using a bioanalyzer and quantified by Qubit high sensitivity assay. All samples were barcoded, pooled, and run on a single sequencing lane of a HiSeq3000 to generate 50 bp single end reads. RNA-seq reads were analyzed for both quality and adapter contamination using FastQC (http://www.bioinformatics.babraham.ac.uk/projects/fastqc/). All reads were trimmed using Trimmomatic (Bolger et al., 2014) using Illuminaclip single end adapter trimming, a removal of the first three bp from the 5’ as recommended for SMARTer libraries, and a 4 bp sliding window trim for quality scores less than 15. All remaining reads greater than 15 bp were then reanalyzed using FASTQC to verify quality. Potential library contamination was examined by aligning 100,000 reads from each sample to genomic sequences from *E. coli*, mouse (mm10, including rRNA and mtDNA), human (hg38), and PhiX using FastQ Screen (http://www.bioinformatics.babraham.ac.uk/projects/fastq_screen/).

### RNA sequencing PCA analysis

Principle Component Analysis (PCA) was conducted on read counts from published cell type specific RNA sequencing (Zhang et al., 2014; Mo et al., 2015) with normalization for library size. Raw Fastq files for each cell type were downloaded from GEO (GSE63137 and GSE52564) and processed as described except for paired end sequencing. PCA analysis was conducted on the top 500 most variably expressed genes across all cell types using custom R scripts. To further assess microglial purity subsampling of the aligned reads for both neuronal and microglial samples(Zhang et al., 2014) (GEO GSE52564) were performed to match the average 3x10^7^ reads/sample (Table S2) obtained in this experiment. Specific proportions of randomly selected reads were then extracted from the neuron and microglial samples, combined, counted and processed in EdgeR with normalization for total library size. Proportions for each combined mixed sample are listed in Figure S1.

### RNA sequencing differential expression analysis

RNA-seq reads were aligned to the current mouse genome mm10 using the splicing-aware aligner Tophat2 (v2.0.14/bowtie2.2.5, stranded, Single End)(Kim et al., 2013) and reads aligning to each gene were counted using FeatureCounts (Liao et al., 2014) with exclusion of multimapping reads. The resulting table of counts per gene and sample was analyzed using the EdgeR pipeline (Robinson et al., 2010). Lowly expressed genes (less than 1 count per million reads in at least three biological replicates from either MAA or PBS groups) were removed from the analysis and the remaining reads were normalized for library size and RNA composition using the calcNormFactors function with trimmed mean of M-size method (Robinson and Oshlack, 2010). The normalized libraries were then fed into a one factor model (MAA vs. PBS) with a zero intercept. Both common and tagwise dispersions were estimated using estimateDisp and the resulting model was used for differential expression using exactTest and topTags with a filter for false discovery rate < 0.05. All differential expression analysis was performed on normalized counts per million (CPM) mapped reads. For graphical comparison of selected genes of interest and heatmap generation Reads per kilobase of transcript per million mapped reads (RPKM) was calculated using the built-in rpkm function that normalized for gene length. Gene Ontology and Pathway enrichment for differentially expressed genes were conducted using Enrichr (http://amp.pharm.mssm.edu/Enrichr/enrich) (Chen et al., 2013; Kuleshov et al., 2016) and then trimmed using REVIGO (http://revigo.irb.hr/) (Supek et al., 2011).

### Whole Genome Bisulifte sequencing (WGBS) analysis

WGBS libraries were constructed using the Illumina EpiGnome/TruSeq DNA Methylation kit (Illumina, EGMK81312) with indexing barcodes (Illumina, EGIDX81312). 100 ng of DNA from each sample was bisulfite converted using the EZ DNA Methylation-Lightning kit (Zymo, D5030). Each library was given a unique barcode identifier and 14 cycles of PCR amplification. Libraries were quantified by Qubit and quality was assessed by Bioanalyzer. Sequencing of barcoded and pooled libraries was performed across two lanes of Illumina HiSeq4000 to obtain 100 bp single end reads.

After adapter trimming and quality assessment, all WGBS reads were mapped to the mouse genome (mm10) using the Bisulfite Seeker2 program (v2.0.8 with bowtie1.1.1) (Guo et al., 2013) and subsequent analysis was done with custom Perl and R scripts (Dunaway et al., 2016b). Differentially methylated regions (DMR) between experimental conditions was examined using the R packages DSS and bsseq (Feng et al., 2014). DMRs were identified as sets of CpGs with a t-statistic greater than the critical value for alpha 0.05 and with a gap between CpGs of less than 300 bases. DMRs were then filtered for those with more than 3 CpGs, a mean methylation difference of greater than or equal to 10% between conditions, and an area stat of +/- 20. Further permutation testing was used to identify DMRs with a family wise error rate <0.05 based on 1,000 permutations of random shuffling of sample group assignment. The average difference in methylation was 29.94% for hyper-methylated and 31.57% for hypo-methylated DMRs. The average size of the hyper-methylated DMRs were 492 bases with an average of 10 CpGs and hypo-methylated DMRs were on average composed of 450 bases and 10.9 CpGs.

DMRs were assessed for Ontology and pathway enrichment using Genomic Regions Enrichment of Annotations Tool (GREAT) with default settings for regulatory regions (5 kb +/- 1 kb of TSS and basal regulatory domain extension from nearest gene of 1Mb) (McLean et al., 2010) with filters for both significant binomial and hypergeometric tests (FDR < 0.05 The resulting GO term lists were trimmed to remove redundancy using REVIGO. Motif enrichment within DMRs was assessed using HOMER (Heinz et al., 2010) (FDR < 0.05) and lists were filtered for motifs commonly identified in both hyper- and hypo-methylated DMRs, hypo-methylated DMRs only, or hyper-methylated DMRs only. Genes with transcription start sites within +/- 5kb of each DMR were identified using HOMER (Heinz et al., 2010).

In addition to global assessment of CpG and CpH methylation levels, CpG methylation over gene bodies, CpG islands, CpG shores (+2 kb from a CpG island), and tissue specific enhancer regions were also examined and tested for differential methylation using independent, two-tailed t-test with FDR correction and a minimum of 5 reads per region across all samples.

### WGBS PCA analysis

PCA was conducted on percent mCG/CG from publish cell type specific WGBS (Lister et al., 2013; Mo et al., 2015). Raw Fastq files for each cell type were downloaded from GEO (GSE63137 and GSE47966) and processed as described for WGBS. Percent CG/CG methylation was extracted for 20 kb windows, gene bodies, and CpG islands for each sample. CpG Islands were masked from the analysis of the other two regions. PCA analysis was conducted on the top 500 most variable regions of each type for all cell types using custom R scripts.

### Fluidigm Biomark HD Delta Gene Expression Assay

Gene expression for selected targets identified from RNAseq analysis were examined in an independent set of microglial samples isolated from PND35 female littermates of animals used for sequencing (Supplemental Table S3). Gene expression analysis was performed using the Fluidigm Biomark HD Delta gene assay on a 48.48 Integrated Fluidic Circuit. Delta gene assay design was performed by Fluidigm and primer sequences are listed in Supplemental Table S3. 20ng of total RNA was used as input for cDNA synthesis (Fluidigm, 100-6472 B1) followed by pre-amplification (13ng per sample) (Fluidigm, 100-5875 C1) using a mix of all forward and reverse primers. Samples were then diluted five-fold and analyzed on a 48.48 IFC and Biomark HD machine (1.4ng/sample) in duplicate (Fluidigm, 100-9791 B1). Cycle threshold values were averaged across replicates and then normalized to *hprt* expression (no difference in mean Ct values for *hprt* between conditions, t= 1.27, p = 0.239) using the delta Ct method (Ct gene – Ct hprt). These values were then used for group comparisons with a two-tailed t-test without an assumption of equal variance. For display purposes, the delta delta Ct values were then calculated relative to the average Ct value of the PBS treated control for each gene, and relative expression levels were then calculated for each group by 2^-delta delta Ct (Figure S4D).

### Overlap of Gene lists

Gene list overlap was performed using the GeneOverlap (Shen, 2013) package in R with a Fisher’s exact test and a False Discovery Rate correction to p=0.05. Gene lists and references are listed in Table S7 (Voineagu et al., 2011a; Oskvig et al., 2012; Garbett et al., 2012; Sugathan et al., 2014; Gilissen et al., 2014; Gunawardhana et al., 2014; Gupta et al., 2014; Iossifov et al., 2014b; Cotney et al., 2015; Cronk et al., 2015; Sanders et al., 2015; Holtman et al., 2015; Matcovitch-Natan et al., 2016; Richetto et al., 2016; Rube et al., 2016; Dunaway et al., 2016b; Zhao et al., 2017; Lombardo et al., 2017). For comparisons to differentially expressed genes the background gene list was all genes identified as expressed in the RNAseq experiment (see above description) (14,301 genes). For comparisons to DMR associated genes, background genes were defined from a call to the DMR finder for mm10 with no filters or cutoffs. This produces a list of all potential locations in the mm10 genome that could be called as a significant DMR. Genes associated with (+/- 5 kb) this background DMR set were used as the background list (22,412 genes). For comparisons between differentially expressed and DMR associated genes, a common background gene set was used that included the intersection of both the differentially expressed and DMR associated background gene list (13,694 genes that could have both been differentially expressed and within 5 kb of a DMR). All gene lists used in each comparison were filtered to include only genes present on the respective background list for that comparison. Drug target analysis was performed using The Drug Gene Interaction Database (dgidb.genome.wustl.edu)(Dunaway et al., 2016b).

### Results

Figure 1A shows the experimental design prior to microglial isolation. To induce a MAA response, female mice were pre-sensitized with OVA exposure at 6 and 7 weeks of age prior to breeding. Timed pregnant dams were then given three exposures to ovalbumin (OVA) during gestation (gestation day 9.5, 12.5, and 17.5) or three exposures to endotoxin free PBS to mimic immune challenge or control, respectively (Schwartzer et al., 2015).

**Figure 1.**
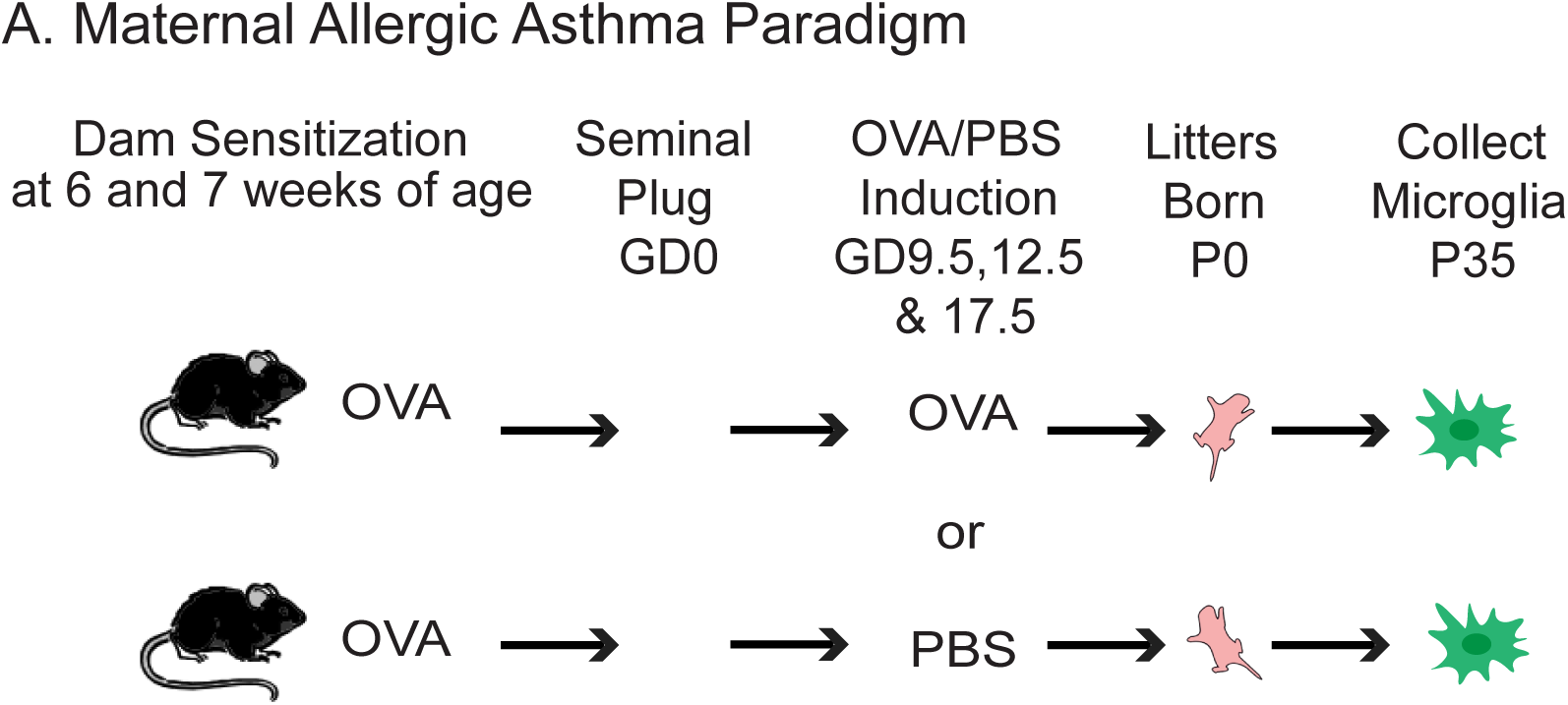
Maternal Allergic Asthma Model. A. Maternal allergic allergy/asthma (MAA) was induced by pre-sensitizing dams to ovalbumin (OVA) prior to mating. During subsequent pregnancy half of the dams were challenged with OVA (gestation day (GD) 9,12, and 17) and the other half received PBS. Microglia were then harvested from the resulting female pups on postnatal day 35 (P35). Male pups were used for behavioral assessment as part of a recently published study (Schwartzer et al., 2017).

While ASD has a pronounced sex-bias for males, both male and female MAA offspring show similar behavioral phenotypes deficits in social interactions, increased marble burying, and decreased grooming (Schwartzer et al., 2015, 2017). In rodents, adolescent females show increased microglial reactivity compared to males (Schwarz et al., 2012). Consequently, this study focused on characterizing juvenile microglia from female MAA offspring. In order to study the effects of *in utero* MAA on persistent microglial epigenomic signatures that coincide with the observance of ASD-like behaviors (Schwartzer et al., 2015, 2017), microglia were isolated from juvenile mice from multiple litters (n= 4 mice from a total of 4 litters per condition) on P35. There were no significant differences in the number of microglia isolated or the RNA/DNA yields between the two treatment groups (Table S1).

## Microglial Methylome and Transcriptome Reveal Cell Type Specific Signatures

RNA sequencing was performed to examine differences in gene expression associated with MAA in juvenile microglia (n=4/group). There were on average 28.6 million uniquely mapped reads per sample and no differences in read depth or mapping between conditions (Table S2). All libraries passed FastQC assessment and none indicated significant sources of contamination during library preparation (Figure S1A & B).

We first sought to ensure that the gene expression patterns in both experimental conditions fit with expected microglial markers and gene expression patterns. To examine cell type specific gene expression patterns we compared the MAA and PBS microglia collected in this experiment to previously published RNA sequencing data on carefully sorted cell populations from mouse cerebral cortex (neurons, astrocytes, oligodendrocyte precursor cells, newly formed oligodendrocytes, myelinating oligodendrocytes, microglia, endothelial cells, and pericytes) (Zhang et al., 2014) and several specific types of neurons (excitatory, PV interneurons and VIP interneurons) (Mo et al., 2015). Both the PBS and MAA microglia transcriptomes cluster more closely with those of adult cortical microglia and are separated from those of other cortical cell types (Figure 2A). Further analysis of expression (reads per kilobase of transcript per million mapped reads, RPKM) values for cell type marker genes (Hickman et al., 2013; Zhang et al., 2014) revealed high expression of microglial markers and low expression of cell type specific markers from other cortical cell types (Figure 2B). In addition, subsampling microglial and neuronal reads from previously published RNAseq (Zhang et al., 2014) indicated that the percentage of neuronal RNA contamination in our microglial samples was minimal and between 0.2 to 4% (Figure S1C). Analysis by flow cytometry on samples prepared in parallel indicate a similar purity greater than 90% (Figure S1D).

**Figure 2.**
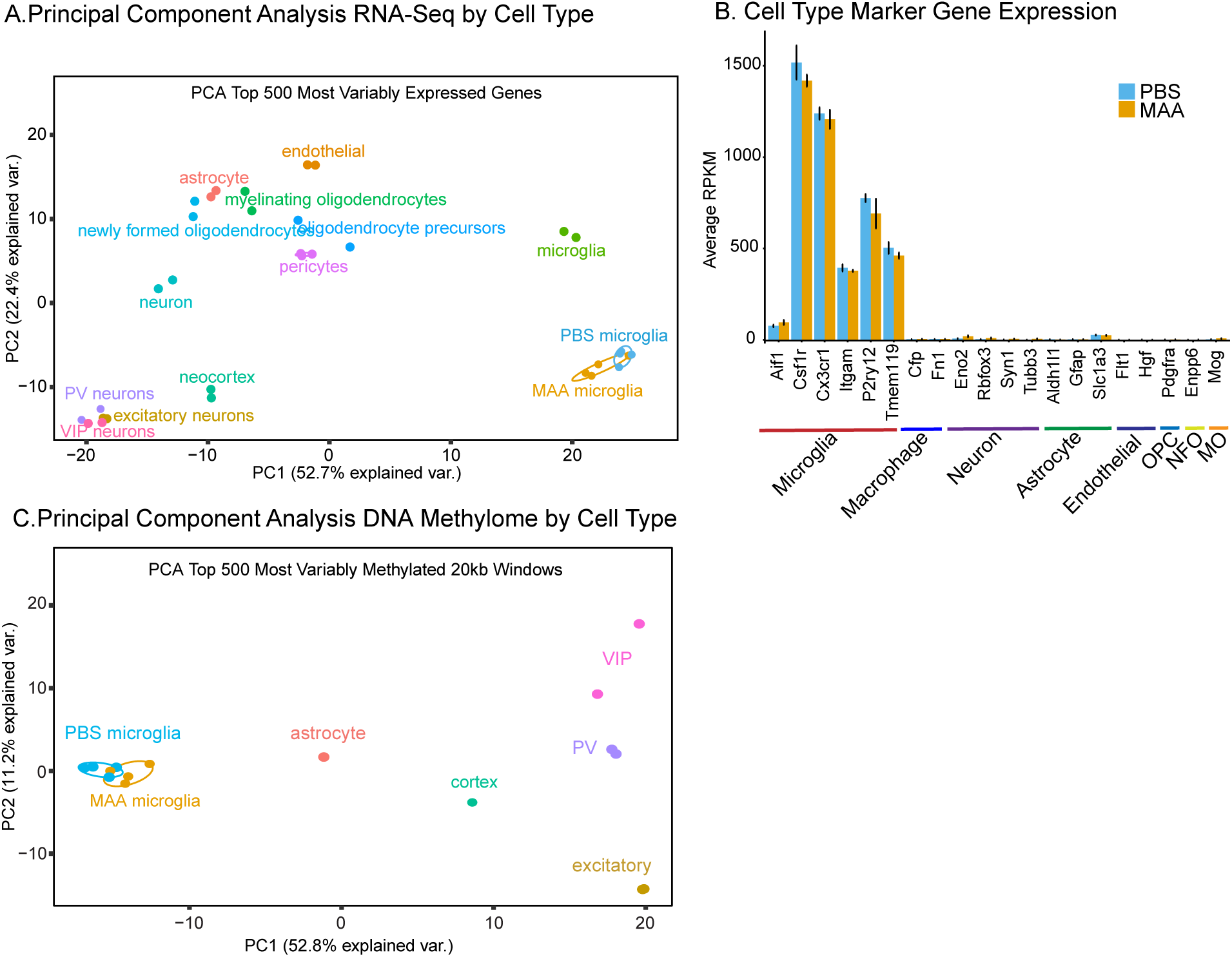
Microglial Transcriptome and Methylome Reveal Cell Type Specific Signatures. A. Principle component analysis of the gene counts from the top 500 most variably expressed genes across multiple brain cell types (neurons, astrocytes, oligodendrocyte precursor cells, newly formed oligodendrocytes, myelinating oligodendrocytes, microglia, endothelial cells, and pericytes (Zhang et al., 2014) and several specific types of neurons (excitatory, PV interneurons and VIP interneurons (Mo et al., 2015)). B. RPKM values for selected cell type marker genes (Hickman et al., 2013; Zhang et al., 2014) for MAA and PBS samples (n=4/group). Oligodendrocyte precursor cells (OPC), newly formed oligodendrocyte (NFO), myelinating oligodendrocyte (MO). Error bars +/- SEM C. Log2RPKM values for MAA compared to PBS microglial RNAseq samples (n=4/group). C. Principle component analysis of the %mCG/CG from the top 500 most variably methylated 20 kb windows across the genome of multiple brain cell types including several specific types of neurons (excitatory, PV interneurons and VIP interneurons (Mo et al., 2015)), astrocytes and total fetal cortex (P28) (Lister et al., 2013). CpG Islands were masked from the analysis.

To examine DNA methylation changes in P35 microglia between MAA and PBS treated samples using an unbiased genome-wide approach, we performed WGBS. There were no significant differences in sequencing coverage, bisulfite conversion, or in the global percentage of CG, CHG, or CHH methylation between treatments (Table S2). On average microglia showed similar levels of mCG and mCH as previously described in neurons (Lister et al., 2009). To examine cell type specific methylation patterns we compared 20 kb windows across the genome from the MAA and PBS microglia collected in this experiment to previously published WGBS data on several specific types of neurons (excitatory, PV interneurons and VIP interneurons) (Mo et al., 2015), astrocytes, and brain (P28) (Lister et al., 2013). Both the PBS and MAA P35 microglia samples cluster more closely with each other than with other cortical cell types, indicating microglia show a distinct methylation pattern from other brain cell types (Figure 2C). Similar patterns were observed over gene bodies and to a lesser extent over CpG islands (Figures S2A and B). There were no significant differences between MAA and PBS treatment groups in average percent CG methylation for a number of global genomic features, including gene bodies, 5 kb upstream promoter regions (with or without CpG islands), nor CpG islands (Figure S2C).

## Differentially Methylated Regions in MAA Offspring Microglia

To assess the long-term impact of MAA on epigenetic regulation in offspring microglia we identified locus-specific differentially methylated regions (DMRs) between MAA and PBS (Dunaway et al., 2016a; b). 1,184 DMRs were identified, with nearly equivalent numbers of hyper-methylated (626 MAA>PBS) and hypo-methylated (558 PBS>MAA) regions (Figure 3A, Table S3). Three DMRs also passed family wise error rate (FWER) correction (Figure S2D). In general, MAA associated DMRs were largely localized within introns (40%) and intergenic sites (26%) (Figure 3B) indicating potential regulatory function.

**Figure 3.**
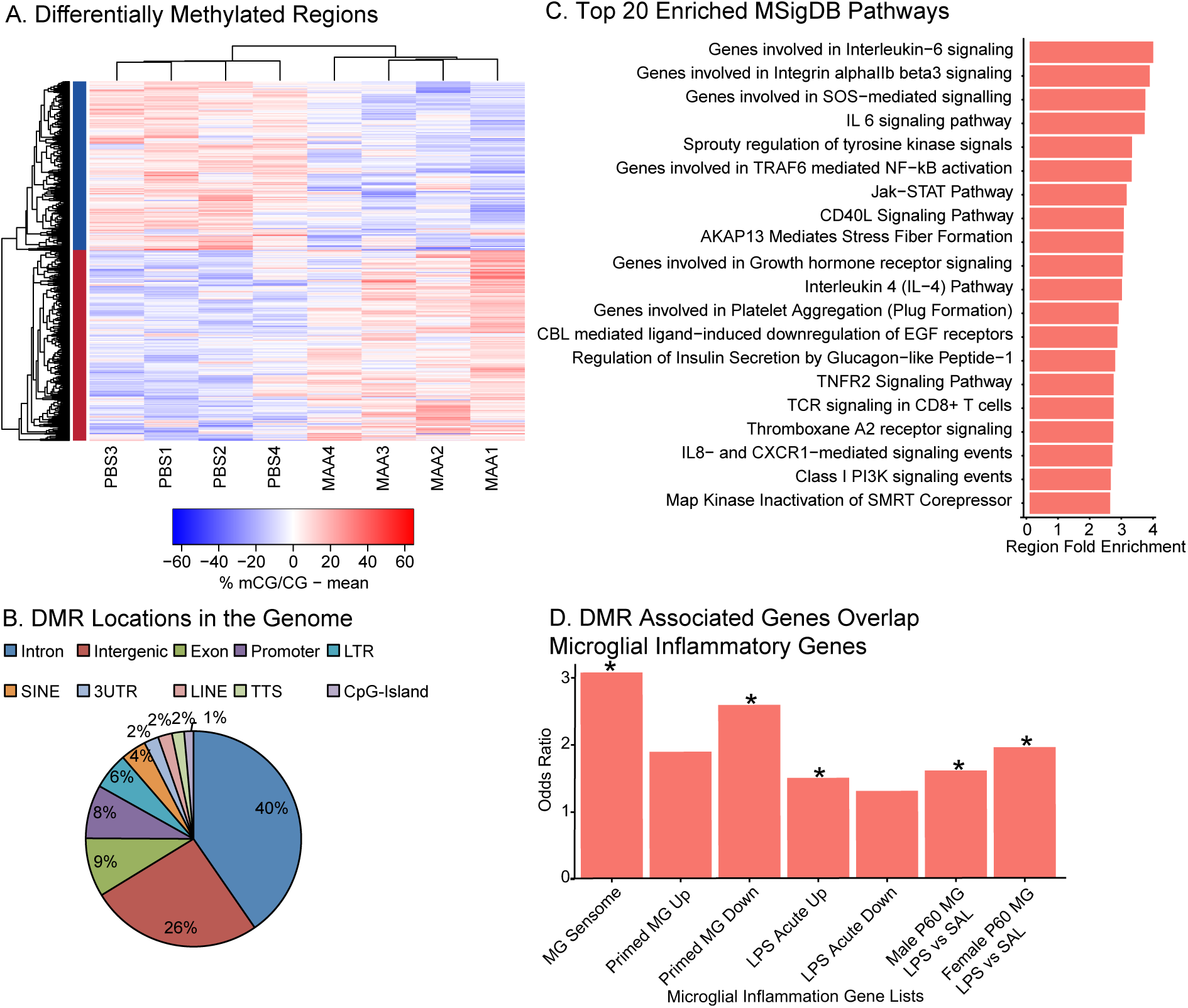
Differential Methylation between MAA and PBS Microglia. A. Differentially methylated regions (DMRs) with an area statistic >= +/-20 and a 10% or greater difference in average methylation between treatments. 1184 DMRs were identified in two clusters: hyper-methylation (MAA>PBS, 626 and hypomethylation (PBS>MAA, 558). Values shown in the heatmap are the difference from the mean for mCG methylation: % mCG/CG minus the mean of %mCG/CG for all samples. See Table S3. B. Distribution of DMRs relative to Transcription Start Sites within +/- 5kb of the DMR (HOMER (Heinz et al., 2010) annotation). C. Top 20 significantly enriched pathways from MSigDB using GREAT annotation of 1184 DMRs. Region Fold Enrichment: observed number of DMRs with the annotation/expected number of DMRs with the annotation (McLean et al., 2010). See Table S4 for full lists. D. Fisher’s Exact Test Odds Ratios for gene list overlap between genes with transcription start sites +/- 5kb from the MAA DMRs. DMR associated genes significantly overlapped genes from the microglial (MG) sensome (Hickman et al., 2013), primed microglia, genes up-regulated acutely in response to LPS (Holtman et al., 2015), and genes up-regulated in microglia two hours after LPS in males and females at postnatal day 60 (P60) (Hanamsagar et al., 2017). See Table S7 for full lists and statistics.

Functional annotation was assigned to each DMR using GREAT, which included biological processes such as regulation of cell number, embryo development, and response to cytokine stimulus. Pathway analysis (Panther and MsigDB) showed enrichment for a number of immune pathways including inflammation mediated by cytokine and chemokine signaling such as IL-6, IL-4 and IL-8 signaling, as well as Jak-STAT, NF-kB, TNF and mTOR signaling (Figure 3C and Table S3). When specific enrichment was examined for MSigDB immune signatures, DMRs were found to be enriched for gene sets associated with LPS treatment and other types of immune cell activators (Table S3). Furthermore, genes with transcription start sites +/-5kb of DMRs (424 DMRs, total with 209 hyper-methylated DMRs and 215 hypo-methylated DMRs) (Table S5) were significantly enriched for genes responsive to LPS, and other immune stimuli in microglia (Figure 3D) (Hickman et al., 2013; Holtman et al., 2015; Hanamsagar et al., 2017). Together these findings indicate that regions sensitive to MAA methylation differences may contain critical regulatory regions responsive to immune stimuli in microglia.

To evaluate potential transcription factor binding sites enriched within MAA-DMRs, we examined motif enrichment analysis using HOMER (Table S4). Motifs commonly enriched in hypo- and hyper-methylated DMRs include transcription factors related to a number of different functions including the SOX, SMAD, and TGFβ-induced families of transcription factors.

Hyper-methylated DMRs were enriched for several interesting transcription factor motifs that are critical for early microglial development and immune activation including RUNX1, PU.1, IRF8, NF-kB, and MAFB. Hypo-methylated DMRs were enriched for several zinc finger and LIM homeobox proteins, together suggesting that regulation of inflammatory signaling and microglial developmental may be altered by DNA methylation differences in MAA microglia.

## Differentially Expressed Genes in MAA Offspring Microglia

In order to understand how epigenomic differences in MAA offspring microglia were related to regulation of gene expression we examined differential gene expression patterns from the same microglial isolations that were profiled for DNA methylation. Differential expression between MAA and PBS samples was conducted using the standard EdgeR pipeline and revealed 162 differentially expressed genes at an FDR < 0.05 (Figure 4A & B and Table S5). Of these genes the vast majority showed greater expression in MAA samples compared to PBS (157 MAA>PBS). Gene Ontology enrichment analysis revealed significant enrichment of several categories of GO terms including gated and ion channel activity, synapse part, and regulation of cell projection organization (Figure S3A and Table S6). Pathway enrichment using KEGG, Panther and Reactome revealed significant enrichment for pathways involved in a number of processes critical for neurodevelopment including several neurotransmitter systems, axon guidance, and synapse function (Table S6). While many of these gene targets are typically examined in neurons, microglia express GABA_B_, serotonin and dopamine, NMDA, AMAP, and kainic acid and metabotropic glutamate receptors that can be utilized to sample the local environment and consequently regulate activation, chemotaxis, secretion, proliferation and phagocytosis (Craner et al., 2005; Li et al., 2008; Black and Waxman, 2012; Stebbing et al., 2015). Gene expression changes were further validated on an independent group of microglial samples using a Fluidigm Delta Gene Expression Assay on a selected subset of transcripts (Table S3B). With this independent method and cohort all transcripts validated the direction of expression change seen in the sequencing cohort (Table S5) with *Hspalb* levels reaching significance (p < 0.05) and *Ina* levels approaching significant (p < 0.07) (Figure S3B).

**Figure 4.**
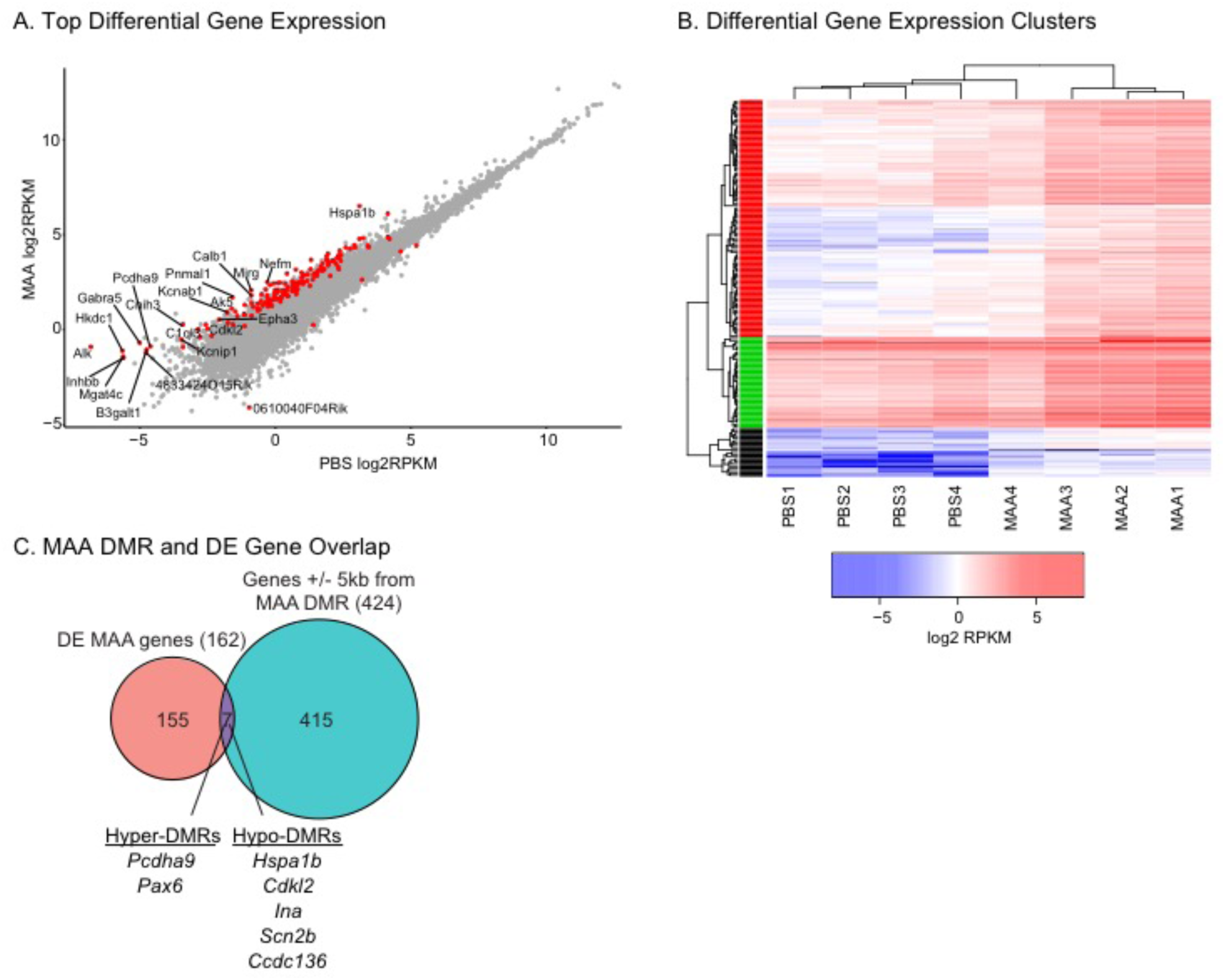
RNA Sequencing Analysis of P35 Microglia. A. Significantly differentially expressed genes (162; FDR q <0.05) are highlighted in red. Genes with log fold change >= 2.5 are labeled. B. Heatmap of log2 RPKM values for the 162 differentially expressed genes between MAA and PBS conditions. Hierarchical clusters (k=3) were used to identify patterns of gene expression using hclust in R. See Table S5 for gene lists, RPKM values, and statistics. C. MAA differentially expressed and DMR associated (+/- 5kb from a DMR) share seven genes. See Table S7 for statistics.

To investigate the relationship between MAA-induced methylation and expression changes, DMR associated genes were compared with differentially expressed genes. There was not a significant overlap between all differentially expressed genes and all genes associated with DMRs (Fisher’s exact test, Odds Ratio = 1.96, p = 0.116) with only 7 genes shared between the two lists (Figure 4C). There was also not a significant overlap with the 70 genes identified as associated with single site CpG differential methylation from blood from human infants from asthmatic mothers (Fisher’s exact test, all p > 0.05) (Gunawardhana et al., 2014). For the seven genes that did overlap between MAA differentially expressed and DMR associated genes, *Pcdha9*, *Pax6* and *Ina* are all critical genes for normal neurodevelopment and are SFARI ASD risk genes (SFARI.org), suggesting that alterations in MAA microglial gene expression and methylation may impact similar pathways disrupted by ASD genetic mutations.

## MAA Microglial Impacted Genes are Enriched for Genes Dysregulated in ASD

In order to more fully evaluate the impact of MAA on ASD-related risk genes we overlapped the differentially expressed and differentially methylated gene lists with previously published risk gene lists for ASD, schizophrenia, and intellectual disability (ID) (Sanders et al., 2012, 2015; Gilissen et al., 2014; Iossifov et al., 2014a) (Figure 5A and Table S7). Genes that were differentially expressed and genes that were associated with DMRs in MAA microglia significantly overlapped a hand curated list of SFARI ASD risk genes and genes with Likely Gene Disrupting (LGD) mutations in exome-sequencing from ASD probands (Iossifov et al., 2014). However, the majority of known genetic risk factors were not found within the MAA microglial gene lists, suggesting that MAA is largely targeting gene pathways independently impacted by genetic mutations linked to ASD and ID. This is not surprising given that the majority of genetic risk factors for ASD and ID are not immune-related (Needleman and Mcallister, 2012; Sanders et al., 2015). However, differences in gene expression profiling from human ASD brain have revealed changes in expression of immune signaling pathways and microglial genes (Voineagu et al., 2011; Gupta et al., 2014; Parikshak et al., 2016). To test the hypothesis that MAA microglia may show similar alterations in gene expression as human ASD brain samples, we overlapped the MAA microglial dysregulated genes with genes that showed altered expression or differential epigenomic marks (DNA methylation or histone acetylation) in human ASD brain samples (Voineagu et al., 2011; Gupta et al., 2014; Dunaway et al., 2016b; Parikshak et al., 2016; Sun et al., 2016) (Figure 5B). Differentially expressed genes in MAA microglia significantly overlapped genes with lower expression in both idiopathic and genetic (15q11-13 duplication) forms of ASD as well as genes with increased acetylation. Furthermore, both MAA differentially expressed and DMR associated genes showed enrichment in a number of modules identified from network analysis of differential ASD gene expression from human ASD cortex (Gupta et al., 2014; Parikshak et al., 2016) (Figure 5C and Table S7), suggesting some common pathways are disrupted in human ASD brain and MAA microglia.

**Figure 5.**
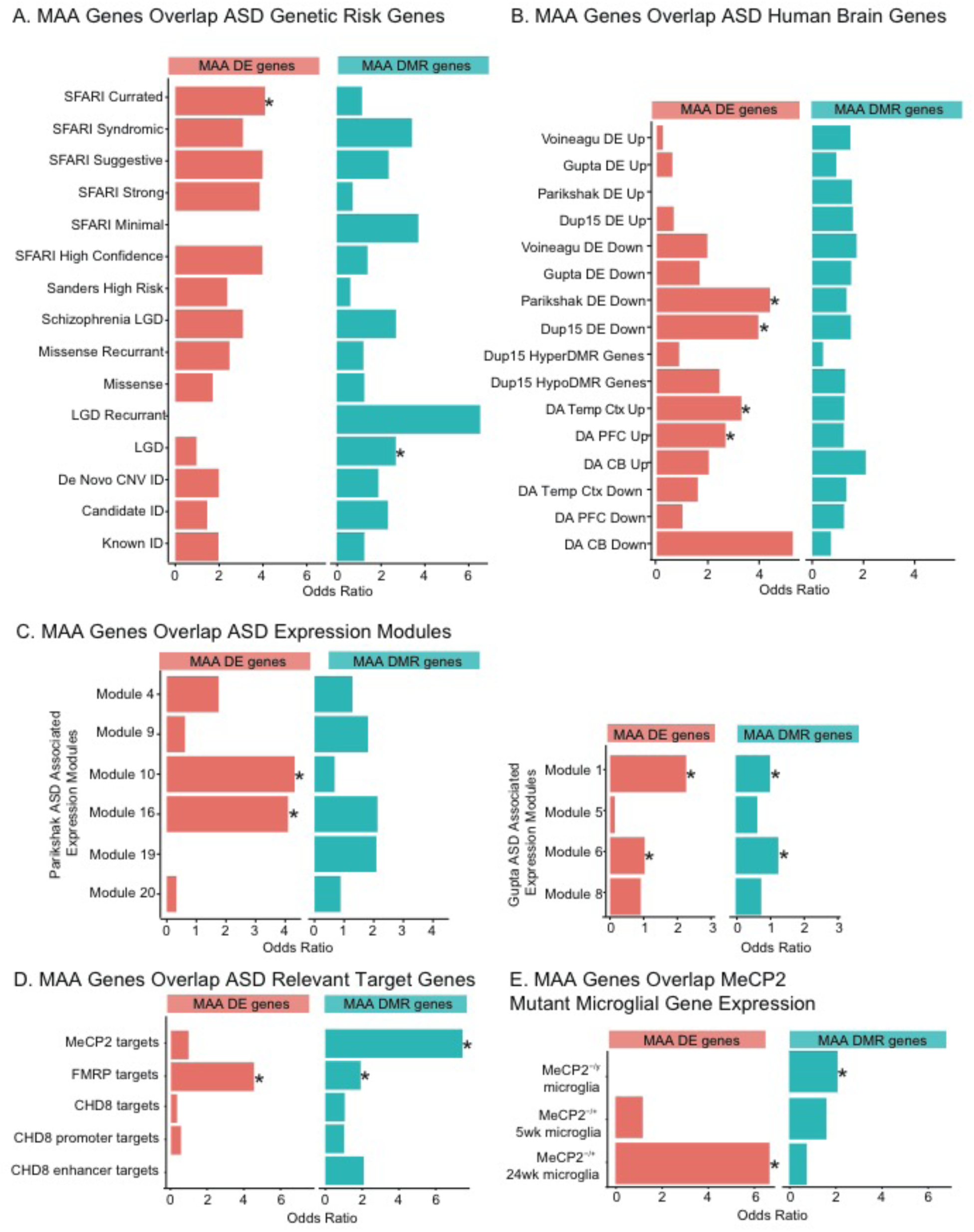
MAA Differentially Expressed and DMR Associated Genes Overlap Genes. Dysregulated in ASD. A. Gene list overlaps between MAA differentially expressed genes (MAA DE genes), MAA DMR associated genes (MAA DMR genes) and ASD genetic risk genes (Sanders et al., 2012, 2015; Gilissen et al., 2014; Iossifov et al., 2014a). C. Gene list overlaps between MAA DE genes, MAA DMR associated genes and genes differentially expressed, methylated, or acetylated in human ASD brain samples (Voineagu et al., 2011; Gupta et al., 2014; Dunaway et al., 2016b; Parikshak et al., 2016; Sun et al., 2016). D. Gene list overlaps between MAA DE genes, MAA DMR associated genes and MeCP2, FMRP, or CHD8 target genes (Darnell et al., 2011; Sugathan et al., 2014; Cotney et al., 2015; Rube et al., 2016). E. Gene list overlaps between MAA DE genes, MAA DMR associated genes and genes differentially expressed in microglia from Rett syndrome mouse models (Cronk et al., 2015; Zhao et al., 2017). Lists are from MeCP2^-/y^ null males (fully symptomatic), MeCP2^-/+^ heterozygous females at either 5 weeks (prior to symptom onset) or at 24 weeks (fully symptomatic). * Fisher’s exact test with FDR corrected p-value <0.05. Gene lists, citations and individual statistics are listed in Table S7.

The enrichment of dysregulated genes in MAA microglia with transcriptionally and epigenetically dysregulated genes in ASD brain suggests that MAA may alter similar pathways as those impacted by ASD. Consequently, we examined the overlap between MAA regulated genes and gene targets of three of the most commonly mutated regulatory factors implicated in ASD and ID: Fragile X Mental Retardation Protein (FMRP), Chromatin-Helicase DNA Binding Protein 8 (CHD8), and Methyl CpG Binding Protein 2 (MeCP2) (Figure 4D). FMRP targets from mouse brain (Darnell et al., 2011) overlapped with both MAA differentially expressed and DMR associated genes (Figure 5D and Table S7) suggesting that key gene targets downstream of FMRP may also be impacted by MAA. Similar overlap between FMRP targets and genes acutely regulated 4 hours following LPS induced MIA has also been previously observed (Lombardo et al., 2017), suggesting that FMRP target genes may be specifically vulnerable to maternal inflammation. There was no significant overlap between either MAA gene list and identified CHD8 target genes from multiple datasets (Sugathan et al., 2014; Cotney et al., 2015). MAA DMR associated genes were enriched in MeCP2 target genes identified from high resolution ChIP-sequencing from olfactory epithelium neurons (Rube et al., 2016), suggesting MeCP2 may be associated with sites of MAA differential methylation. MeCP2 has previously been implicated in microglial function in Rett syndrome (Derecki et al., 2012; Cronk et al., 2015; Schafer and Stevens, 2015), and microglia from Rett mouse models show altered gene expression patterns related to glucocorticoid signaling and stress response (Cronk et al., 2015; Zhao et al., 2017). Genes differentially expressed in Rett syndrome mouse model microglia overlapped both MAA differentially expressed and DMR associated genes (Figure 5E and Table S7), further supporting the role for MeCP2 as a reader of DNA methylation in microglia and indicating that MeCP2 function may be negatively impacted in MAA microglia.

## MAA Offspring Microglia Compared to Other MIA Models

Maternal immune activation has been utilized as a model of ASD as well as schizophrenia and general neurodevelopmental impairment (Hagberg et al., 2012; Mattei et al., 2017). The MAA model was originally designed to examine allergic asthma as a risk factor for ASD, but the model may be applicable to other neurodevelopmental disorders that share maternal immune activation as a risk factor. To compare our MAA model to other MIA models we overlapped the MAA gene lists with published datasets from maternal exposures to LPS, PolyI:C, influenza, or direct cytokine injection. DNA methylation in MAA microglia and whole prefrontal cortex from PolyI:C induced MIA (Richetto et al., 2016) showed similar patterns of differential methylation (Figure S4A) suggesting the long-lasting DNA methylation signatures in the brain may be target similar pathways across MIA/MAA models. In comparison, our MAA differentially expressed and DMR associated microglial gene lists largely did not overlap with whole brain gene expression patterns induced four hours following different types of MIA (LPS, recombinant IL6, PolyI:C) (Figure S4B and Table S7). The one exception to this was a significant overlap with genes acutely downregulated in response to LPS MIA. There were mixed findings for gene overlap with microglia isolated from PolyI:C MIA models. MAA microglial gene lists did not overlap dysregulated genes identified in a PolyI:C MIA model from isolated microglia from whole brain on P0 or P28 (Matcovitch-Natan et al., 2016) (Figure S4B). However, MAA differentially expressed genes significantly overlapped with genes that were up-regulated in a PolyI:C MIA model from hippocampal microglia from adult animals (Mattei et al., 2017) (Figure S4C) suggesting that MAA may impact hippocampal microglia. Interestingly, both MAA differentially expressed and DMR associated gene lists overlapped genes that were responsive to minocycline treatment in adult MIA animals (Mattei et al., 2017) (Figure S4C), indicating that minocycline may serve as a potential therapeutic for reversing some of the MAA offspring phenotypes.

### Discussion

In this study we identified epigenetic changes in microglia from juvenile offspring of MAA dams that advance the understanding of maternal immune activation and ASD risk in several important ways. We found differentially methylation regions enriched for transcription factor binding sites related to immune signaling, microglial inflammation, and regulation of microglial development. This immune related methylome signature in MAA offspring microglia implicates altered epigenomic regulation of microglial inflammatory responses as a result of MAA. Importantly for ASD relevance, transcriptome profiling of MAA microglia identified key developmental genes related to synaptic function that are also misregulated in human ASD brain.

The gestational timing of MAA may preferentially affect changes in microglial function in offspring, a perturbation that may be marked by changes in DNA methylation that persist as microglia develop. Microglia migration to the brain and colonization of the neural folds during embryogenesis with minimal turnover (Squarzoni, 2015), suggesting that perturbations to microglia in early development may be maintained throughout the lifetime of the animal. Previous work with PolyI:C MIA demonstrated long-lasting alterations in histone modifications (Tang et al., 2013) and DNA methylation (Basil et al., 2014; Richetto et al., 2014, 2016) in adult brain that stem from in utero exposures at mid gestation. While it is unclear if the methylation changes in juvenile MAA microglia are direct or secondary impacts from MAA, the enrichment for numerous immune related pathways including IL-6, IL-4, and IL-8 signaling, NF-kB activation and JAK/STAT signaling, suggests that DMRs may either serve as a signature of prior immune activation during gestation or as epigenetic mechanisms for priming future immune function. The enrichment of DMRs for IL-4 signaling is consistent with the cytokine profile of elevated IL-4 and IL-5 observed in asthma (Magnan et al., 2000; Cho et al., 2005; Tamasi et al., 2005; Kumar et al., 2006), the mid-gestation serum cytokines profiles observed in women who gave birth to a child later diagnosed with ASD (Goines et al., 2011; Jones et al., 2017), and in amniotic fluid taken from children later diagnosed with ASD (Abdallah et al., 2012). Similarly, the enrichment of DMRs for a number of transcription factor motifs critical for early development of microglia (RUNX1, PU.1, and IRF8) indicate that MAA may shift the developmental regulation of microglial function. For example, RUNX1 is expressed in microglial yolk sac progenitors and plays a critical role in regulating microglial proliferation in early brain development (Ginhoux et al., 2010) by ensuring that developing microglia transition from an amoeboid to mature, ramified state during differentiation (Zusso et al., 2012). Similarly, PU.1 is a pioneer transcription factor with DNA methylation sensitive binding (Stephens and Poon, 2016) critical for microglia progenitor development (Kierdorf et al., 2013; Goldmann et al., 2016), regulates microglia specific enhancers (Gosselin et al., 2014) and gene expression (Satoh et al., 2014). PU.1 and shows increased expression in human ASD brain(Gupta et al., 2014), and manipulations of PU.1 alter the expression of key microglial genes and phagocytosis function (Huang et al., 2017), indicating that changes in PU.1 binding could dramatically alter microglial function in ASD. Future studies will be needed to examine the impact of MAA on microglia from different stages of development and in altering the transcriptional response to secondary immune activation (i.e. LPS).

In human ASD brain samples, whole genome analysis has identified alterations in DNA methylation (Nardone et al., 2014; Dunaway et al., 2016b) and gene expression (Voineagu et al., 2011; Gupta et al., 2014; Werling et al., 2016) of a number of immune related genes including components of the complement cascade *C1Q*, *C3*, *ITGB2* and the cytokine *TNF*α. Several studies have identified alterations in microglia morphology and density in post-mortem brain samples from ASD patients (Vargas et al., 2005; Morgan et al., 2010, 2012, 2014; Tetreault et al., 2012). In at least a subset of ASD cases, microglia adopt an immune activated morphology characterized by retraction and thickening of processes and enlargement of the soma in multiple brain regions including the dorsolateral prefrontal cortex (Morgan et al., 2010), amygdala (Morgan et al., 2014), and cerebellum (Vargas et al., 2005). Using positron emission tomography and a radiotracer for microglia, Suzuki and colleagues found evidence for increased microglia activation in multiple brain regions in young adults with ASD (Suzuki et al., 2013). Given that immune system related genes are not highly prevalent among the genetic candidates for ASD risk, the alterations in expression and methylation of immune genes in ASD may occur in reaction to abnormal neurodevelopment and/or in response to environmental factors such as maternal immune activation.

The significant overlap of MAA differentially expressed genes with genes differentially expressed or differentially acetylated in ASD brain provides support for the role of maternal immune dysregulation in the etiology of ASD. MAA may also impact some of the same downstream transcriptional pathways that are regulated by MeCP2 and FMRP. While FRMP was not a direct target of MAA in our data, *Cytoplasmic FMRP Interaction Protein 2* (*Cyfip2*) had an intronic hyper-methylated DMR in the MAA microglia. *CYIFP2* expression is altered in Fragile X-syndrome (Hoeffer et al., 2012) and defective CYFIP/FMRP interactions have been implicated in ASD and ID (Abekhoukh and Bardoni, 2014) and may mediate some of the overlapping genes targeted by FMRP and regulated by MAA. MeCP2 expression was also not altered by MAA, but genes targeted by MeCP2 binding and genes sensitive to loss of MeCP2 in microglia significantly overlapped MAA genes, indicating MeCP2 may play a role in regulating gene expression differences in MAA microglia.

The vast majority of differentially expressed genes in MAA microglia showed a significant increase in expression relative to PBS controls. While not expected, this increase in expression is consistent with other MIA experiments in whole brain examining acute responses to MIA treatments (Garbett et al., 2012), suggesting an increased sensitivity to environmental signals. For example, we found increased expression of voltage-gated calcium channel subunit *Cacna1b*, voltage-gated potassium channel subunits (*Kcnab1* and *Kcnd2),* and voltage-gated sodium channel subunits (*Scn1a*, *Scn2a*, and *Scn2b*). *Canca1b*, *Kcnd2* and *Scn1a* are also ASD candidate genes and increased expression of a number of ligand-gated ion channels upon MAA could lead to altered function (Eder, 1998; Craner et al., 2005; Schilling and Eder, 2007, 2015; Li et al., 2008; Black et al., 2009; Wu et al., 2009; Black and Waxman, 2012; Stebbing et al., 2015). Increased surface receptor expression in MAA microglia would be consistent with motif enrichment of DMRs for transcription factors related to adult homeostatic function in microglia (FOS, MEF2A, MAFB)(Matcovitch-Natan et al., 2016). For example, MAFB expression is induced during microglial maturation, its motif is enriched in macrophage-specific enhancers, and deletion of *Mafb* specifically in microglia increases gene expression for immune and viral response genes (Matcovitch-Natan et al., 2016). Together these findings suggest that MAA results in long-lasting changes in the ability of microglia to sense and respond to changes in the local brain environment. Altered sensitivity of MAA microglia could potentially dramatically alter the key role of microglia in sculpting neuronal connections and neuronal cell number in the developing brain (Tremblay et al., 2010; Paolicelli et al., 2011; Hagberg et al., 2012; Schafer et al., 2012; Zhan et al., 2014), as further suggested by enrichment of MAA differentially expressed genes for pathways involved in regulation of cell projection organization, neuronal differentiation, and neuronal apoptotic processes (Table S6). Experimentally removing or manipulating microglia in fetal life or during transient postnatal stages has been demonstrated to have profound effects on the number and strength of neuronal synapses, neuronal circuit development, and behavior (Paolicelli et al., 2011; Cunningham et al., 2013; Zhan et al., 2014; Kim et al., 2016). Previous work showed that early life perinatal inflammation (LPS GD15 and 16) augmented fetal microglia activity and decreased neuronal precursors in the developing rat cerebral cortex (Cunningham et al., 2013), leading to impaired corpus callosum fasciculation (Pont-Lezica et al., 2014), dopaminergic axon outgrowth (Squarzoni et al., 2014), and alterations in glutamatergic synapses in the adult offspring (Roumier et al., 2008). Our results show that MAA microglia have significant transcriptional and epigenetic alterations to genes important in axon guidance and receptor function that are consistent with altered sensitivity to the environment and neuronal circuit development.

One potential limitation of our study was a low level of neuronal contamination (0.2-2% in control versus 0.5-4% in MAA, Fig. S1C) in our microglial isolations due to the inherent microglial-neuronal interactions including phagocytosis that occur in brain. While our microglia isolations were above the purity of another published microglial transcriptome study (Zhao et al., 2017), we cannot be certain that the higher expression of some of the typically neuronal genes were cell intrinsic to microglia rather than a result of increased neuronal phagocytosis in the MAA brain (Solga et al., 2015), although either explanation is of interest for understanding the pathogenesis. While the function of many of these neurotransmitter receptors on microglia is largely unknown (Schafer et al., 2013), their overall increase in expression following MAA suggests that microglia have altered sensitivity in the juvenile MAA brain. This is consistent with recent work from a PolyI:C MIA model that showed altered gene expression of cell surface receptors and phagocytosis activity of adult hippocampal microglia (Mattei et al., 2017). Genes misregulated in this model overlapped the MAA differentially expressed genes indicating shared molecular pathways and functions may be disrupted. In addition, MAA impacted genes also overlapped genes responsive to adult minocycline treatment in this MIA model, suggesting minocycline may be able to similarly reverse the behavioral, gene expression and microglial functional deficits in the MAA offspring. Alternatively, many of the receptors showing increased expression in MAA offspring microglia are identified drug targets for a variety of pharmaceuticals (Table S6) that may provide novel options for therapeutics, as is being explored in other neurologic disorders (Eder, 2010).

This work provides a novel link between maternal allergic asthma and changes in the microglial epigenome, indicating that microglia may serve as a potential therapeutic target for normalizing fetal brain connectivity deficits observed in ASD. Removal of microglia from the adult mouse brain by pharmacological treatment did not negatively impact adult behavior (Elmore et al., 2014, 2015) and improved behavioral deficits in an Alzheimer’s disease model (Rice et al., 2015), suggesting microglial contributions to neuro-immune dysfunction can be reversed in the adult. Given the emerging importance of microglia to multiple neuropsychiatric disorders, uncovering the microglial epigenetic landscape is the first step towards understanding how *in utero* exposures may influence microglia function and dysfunction in later life. Together, our findings present the first whole genome evaluation of microglial DNA methylation and support a role for maternal immune activation in altering microglial function as part of the pathogenesis of ASD.

List of Abbreviations: Ovalbumin (OVA), Maternal Allergy/Asthma (MAA), Phosphate buffered saline (PBS), methylated CpG (mCG), methylated CpA or CpC or CpT (CHH).

## Availability of Data and Material

Processed gene lists and enrichments are given in supplemental tables for each experiment.

## Authors’ Contributions

A.V.C. prepared libraries for sequencing and conducted bioinformatics analysis. M.C. conducted the maternal allergic asthma procedure, collected, and analyzed purity of microglia. A.V.C, J.L. and P.A. wrote the manuscript.

All authors read and approved the final manuscript.

## Compliance with Ethical Standards

The authors declare that they have no conflict of interest.

## Ethical approval

All applicable international, national, and/or institutional guidelines for the care and use of animals were followed. All procedures performed in studies involving animals were in accordance with the ethical standards of the University of California, Davis.

## Acknowledgements

National Institutes of Health T32MH073124-06, 3R01NS081913-11S1, R21HD086669, R21ES025560, IDDRC U54 HD079125 and 5R01NS081913-14, International Rett Syndrome Foundation, Autism Speaks Foundation (#7567), NARSAD foundation, Peter Emch Foundation and the Jane Botsford Johnson Foundation. This work used the Vincent J. Coates Genomics Sequencing Laboratory at UC Berkeley, supported by NIH S10 OD018174 Instrumentation Grant. The fluidigm gene expression experiment was supported by the University of California Davis Flow Cytometry Shared Resource Laboratory with funding from the NCI P30 CA093373, and NIH NCRR C06-RR12088, S10 RR12964 and S10 RR 026825 grants and with technical assistance from Ms. Bridget McLaughlin.

